# Degradation of PET Plastics by Wastewater Bacteria Engineered via Conjugation

**DOI:** 10.1101/2024.02.07.579132

**Authors:** Aaron Yip, Owen D. McArthur, Kalista C. Ho, Marc G. Aucoin, Brian P. Ingalls

## Abstract

Microplastics are contaminants of global concern that pose risks to ecosystems and human health. Focusing on PET plastics, we present a proof-of-concept for reduction of microplastic pollution: *in situ* engineering of bacteria in wastewater to degrade PET. Using a broad-host-range conjugative plasmid, we enabled various bacterial species from a municipal wastewater sample to express FAST-PETase, which was released into the extracellular environment. We found that FAST-PETase purified from some isolates could degrade about 40% of a 0.25 mm thick PET film within four days at 50 °C. We then demonstrate partial degradation of post-consumer PET over 5-7 days by exposure to conditioned media from isolates. These results have broad implications for addressing the global plastic pollution problem by enabling environmental bacteria to degrade PET plastics *in situ*.

## 1 Introduction

Plastics are versatile materials that are lightweight, strong, and chemically-resistant. These desirable properties have made them an integral part of daily activities, infrastructure, and economies since their invention in the 20^th^ century ^1^. Although plastics have been an economic boon—particularly in packaging and construction applications—their use has produced unsustainable levels of waste. From 1950 to 2015, about 4900 megatons, accounting for 59% of all plastics ever produced, have been discarded into landfills and the environment ^2^. Ninety percent of plastics produced—consisting of polyethylene (PE), polypropylene (PP), polyvinyl chloride (PVC), polyethylene terephthalate (PET), polyurethane (PU), and polystyrene (PS) ^2^—are projected to persist for hundreds of years; they are not biodegradable on timescales relative to their end of use ^2^. Consequently, their bioaccumulation poses threats to ecosystems and, potentially, to human health ^4–6^. Annual plastic waste generation is projected to almost triple by 2060 compared to 2019 levels, if trends in current plastic usage continue ^7^. Even with immediate implementation of ambitious strategies for waste management, environmental recovery, and reduction of plastic production, conservative estimates suggest that plastic waste entering terrestrial and aquatic ecosystems will exceed or remain close to 2016 levels between 2030-2040 ^8,9^.

Although PE, PP, PVC, PET, PU, and PS are referred to as non-biodegradable, microbes have ways to metabolize these plastics into their constituent molecules which can then be used as carbon and energy sources ^10–14^. These natural processes are relatively slow, taking weeks to months for significant biodegradation to occur ^13–18^. Biodegradation of some plastics has been accelerated by enhancing the activity and stability of the enzymes involved. For example, PET hydrolase (PETase) from *Ideonella sakaiensis* has been subject to numerous protein engineering efforts that have resulted in PETase variants with depolymerization rates orders of magnitude higher than the wild-type enzyme, and with greater thermo- and chemostability, ^19–23^. The introduction of such engineered enzymes into microbial communities through bioaugmentation ^24,25^ could potentially enhance community plastic-degrading capabilities by a similar level.

Effluent and sludge from wastewater treatment plants are major pathways by which microplastics enter the environment ^26–28^. Secondary treatment processes and digestion of sludge offer opportunities for removal of microplastics through bioaugmentation. As reviewed in ref. 28, microbial communities can be supplemented with engineered bacteria that secrete plastic-degrading enzymes; this can be more cost effective than continuously supplementing a cocktail of purified enzymes into a secondary treatment unit. A major challenge with this cell bioaugmentation approach is persistence of the introduced plastic-degrading microorganisms, which are not adapted to the environment and are therefore likely to be quickly eliminated ^29^. As an alternative strategy that would bypass these barriers to establishment, microbes native to the environment could be genetically engineered to express metabolic pathways for biodegradation of plastic waste. This process, known as genetic bioaugmentation, can be achieved through *in situ* delivery of broad-host-range conjugative plasmids. This genetic bioaugmentation approach has not been used for environmental bioremediation of microplastics, but the technique has been investigated by introduction of catabolic genes into microbial communities for bioremediation of non-plastic-based pollutants in laboratory and pilot scale settings ^30–37^.

Here, we report on the introduction by conjugation of a PETase-coding plasmid into bacteria from municipal wastewater, and demonstrate that these transformed strains can degrade PET plastic. We made use of FAST-PETase, an enzyme variant that is more robust to pH and temperature ranges, and orders of magnitude more efficient than the wild-type enzyme ^21^. We screened for transconjugants and confirmed expression of functional FAST-PETase. We then confirmed direct degradation by conditioned media from some of the transconjugants. Our results highlight the effectiveness of delivering engineered plastic-degrading enzymes to bacteria from wastewater.

## 2 Results

### 2.1 pFAST-PETase-cis conjugates to bacteria from wastewater

We constructed an IncP RK2-based ^39^ conjugative plasmid, pFAST-PETase-cis (Fig. 1a, Supplementary Fig. 1), which carries the FAST-PETase gene ^21^ under control of an arabinose-inducible promoter (P_BAD_) (details in Methods). Our design of pFAST-PETase-cis retains the signal peptide SP_pstu_ and 6×His-tag from the backbone pBTK522::FAST-PETase ^18^ (Fig. 1b). These facilitate secretion and purification of FAST-PETase. We chose to drive FAST-PETase expression from the P_BAD_ promoter to achieve tunable gene expression. Moreover, this promoter has been shown to drive expression in a diverse range of gram-negative bacteria ^40^. The plasmid carries a fluorescent marker (*mCherry*) and two antibiotic resistance markers (Amp^R^, Gm^R^) to facilitate selection and plasmid maintenance.

**Figure 1.**
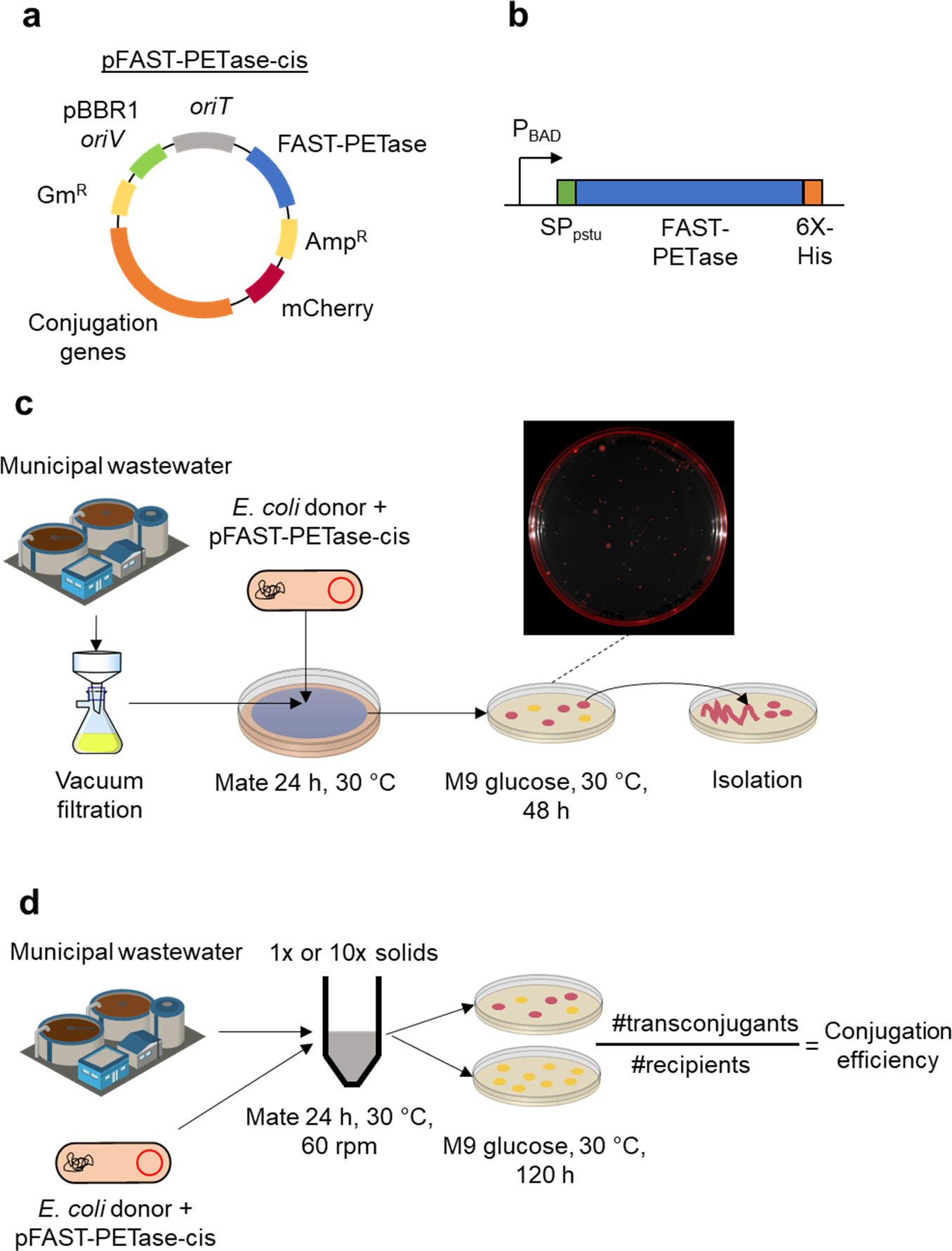
Conjugation of pFAST-PETase-cis into wastewater bacteria. **a** Schematic map of pFAST-PETase-cis (not to scale): *oriT*, RK2/RP4 conjugative origin of transfer; FAST-PETase, gene for FAST-PETase enzyme; Amp^R^, ampicillin resistance gene (*bla-TEM_1_*); *mCherry*, gene for fluorescent protein mCherry; Conjugation genes, encoding the IncP RK2/RP4 conjugation system; Gm^R^, gentamycin resistance gene; pBBR1 *oriV*, plasmid origin of replication. **b** FAST-PETase coding region (not to scale) highlighting the arabinose-inducible promoter (P_BAD_), signal peptide (SP_stu_), and 6×His-tag. **c** Experimental procedure for conjugation of pFAST-PETase-cis into bacteria from a wastewater sample. **d** Experimental procedure for measuring conjugation efficiency in wastewater suspension.

We used *E. coli* K-12 (Δ*ilvD*::*FRT*, Δ*galK*::*cfp-bla,* pSAS31, pTGD, pFAST-PETase-cis) as conjugative donor. This strain is auxotrophic to isoleucine, leucine, and valine, allowing for counter-selection on minimal media (to facilitate isolation of transconjugants). We filter-mated (details in Methods) the donor strain with untreated, municipal wastewater samples from the City of Waterloo, Ontario, Canada, and picked 32 transconjugant colonies expressing *mCherry* on M9 glucose agar plates supplemented with ampicillin and gentamycin (Fig. 1c, details in Methods). No *mCherry* expression was detected in a wastewater sample incubated without donors (Supplementary Fig. 2). Based on bacterial internal transcribed spacer (bITS) sequencing ^41^ on all 32 colonies in a pooled sample (performed by Metagenom Bio Life Science, details in supplementary methods), we observed that most isolates were *Gammaproteobacteria* from the *Enterobacteriaceae* family (Supplementary Data 1), with one exception: *Pusillimonas ssp.*, a gram-negative bacterium from the *Burkholderiaceae* family. From these picked colonies, we selected seven isolates with distinct colony morphologies to identify using bITS sequencing on each individual colony (Table 1, Supplementary Data 2) and to investigate for PET degradation activity.

**Table 1.**
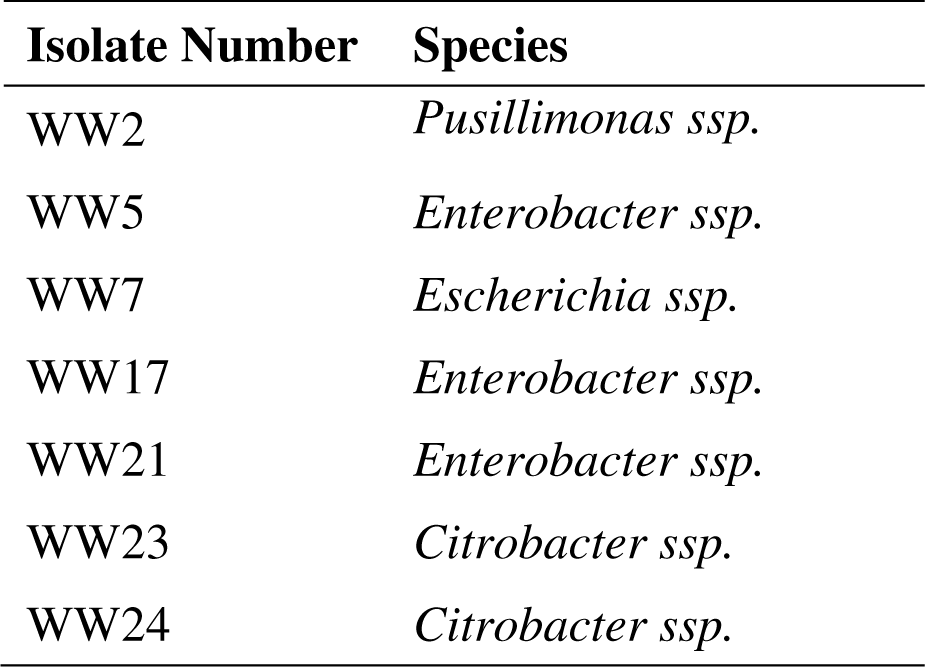
Selected transconjugant isolates identified by bITS sequencing.

To confirm *in situ* delivery in wastewater conditions, we performed a liquid-mating assay where donors were mixed directly into municipal wastewater influent samples (1× or 10× solids) and allowed to mate for 24 h (Fig. 1d). After washing the mating mixture twice in 1× PBS, the mixture was serially diluted and plated onto selective (ampicillin and gentamycin) and non-selective M9 glucose plates, which were incubated for five days at 30 °C. After counting colonies showing *mCherry* activity, we found that the conjugation efficiency (defined as the ratio of transconjugants to transconjugants plus recipients) was on the order of 1% (1.2% with 10x solids, 1.9% with 1x solids). To confirm that observed colonies were not the donor strain, an exponentially growing donor culture was washed twice in 1× PBS and incubated at 30 °C for up to seven days on selective M9 glucose plates—no growth was observed (Supplementary Fig. 2).

### 2.2 pFAST-PETase-cis is lost in absence of selection

To assess the stability of pFAST-PETase-cis, we serially passaged four different transconjugant species once daily in LB media over the course of 14 days. The number of cells expressing ampicillin- and gentamycin-resistance (which are carried on pFAST-PETase-cis) decreased by 4-6 logs for WW2 (*Pusillimonas ssp.*) and WW7 (*Escherichia ssp.*), whereas it decreased by about one log for WW17 (*Enterobacter ssp.*) and WW23 (*Citrobacter ssp.*) (Fig. 2), indicating that the plasmid is lost in the absence of selection.

**Figure 2.**
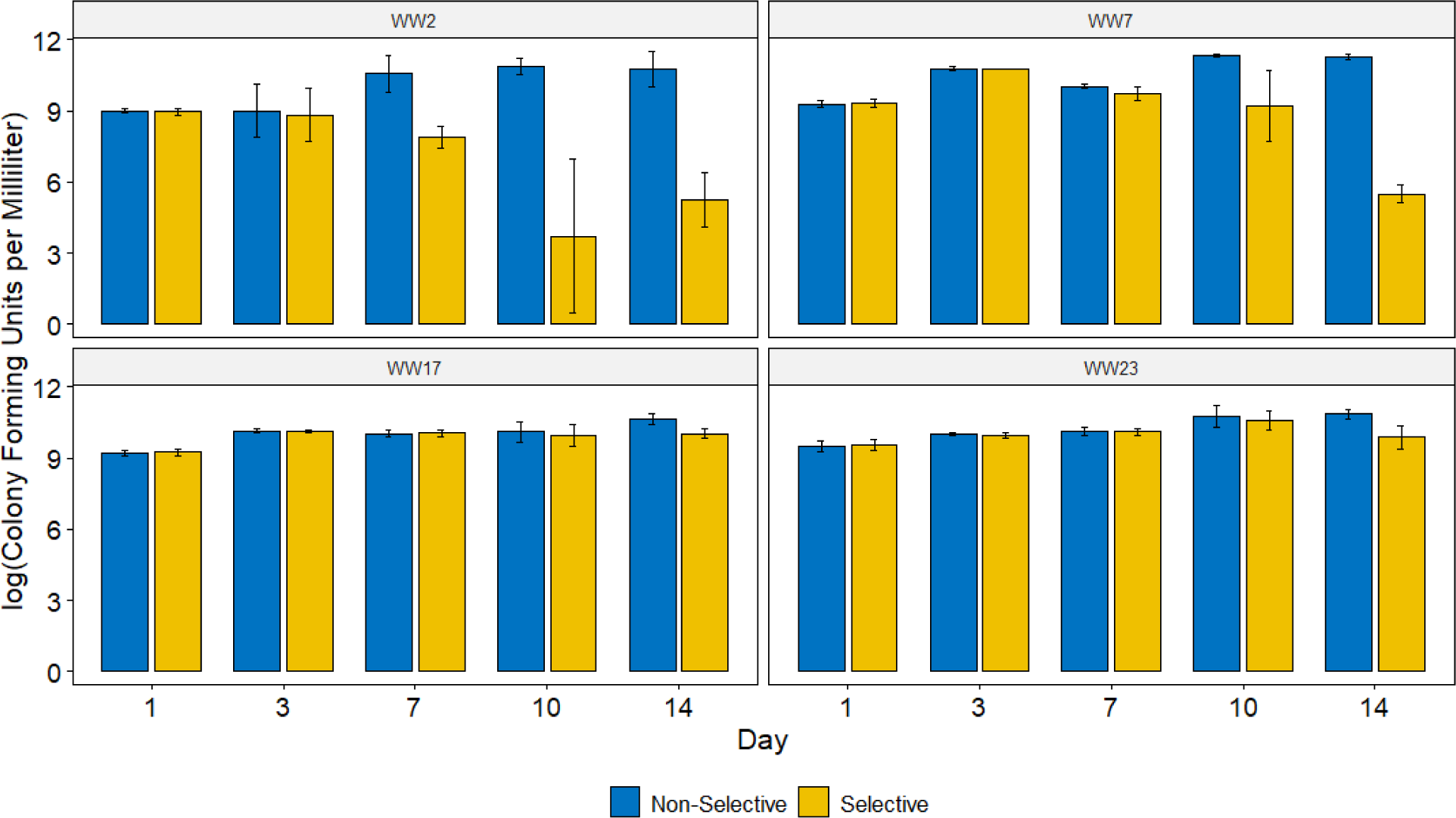
Cell density of wastewater transconjugants plated on selective (ampicillin and gentamycin) and non-selective plates over 14 serial passages in LB. Bars show the mean of three biological replicates. Error bars: ± s.d.

### 2.3 Purified FAST-PETase expressed from transconjugant isolates degrades PET plastic

Before beginning our activity assays, we used SDS-PAGE to characterize intracellular expression of FAST-PETase (Fig. 3a). We then incubated amorphous PET (aPET) film (crystallinity 8.8 ± 1.0% determined by differential scanning calorimetry, Methods) with purified enzyme from cell extracts. Significant mass loss occurred for several isolates (Fig. 3c). We then characterized presence of FAST-PETase in the supernatant by SDS-PAGE (Fig. 3b) and incubated an aPET film with purified enzyme from culture supernatant. Again, significant mass loss occurred (Fig. 3d). Inspection of aPET film samples with scanning electron microscopy showed that samples treated with FAST-PETase (Fig. 3f) had a pitted surface, which is characteristic of enzymatic degradation ^13,21^. In contrast, an untreated aPET sample had a smooth surface (Fig. 3e).

**Figure 3.**
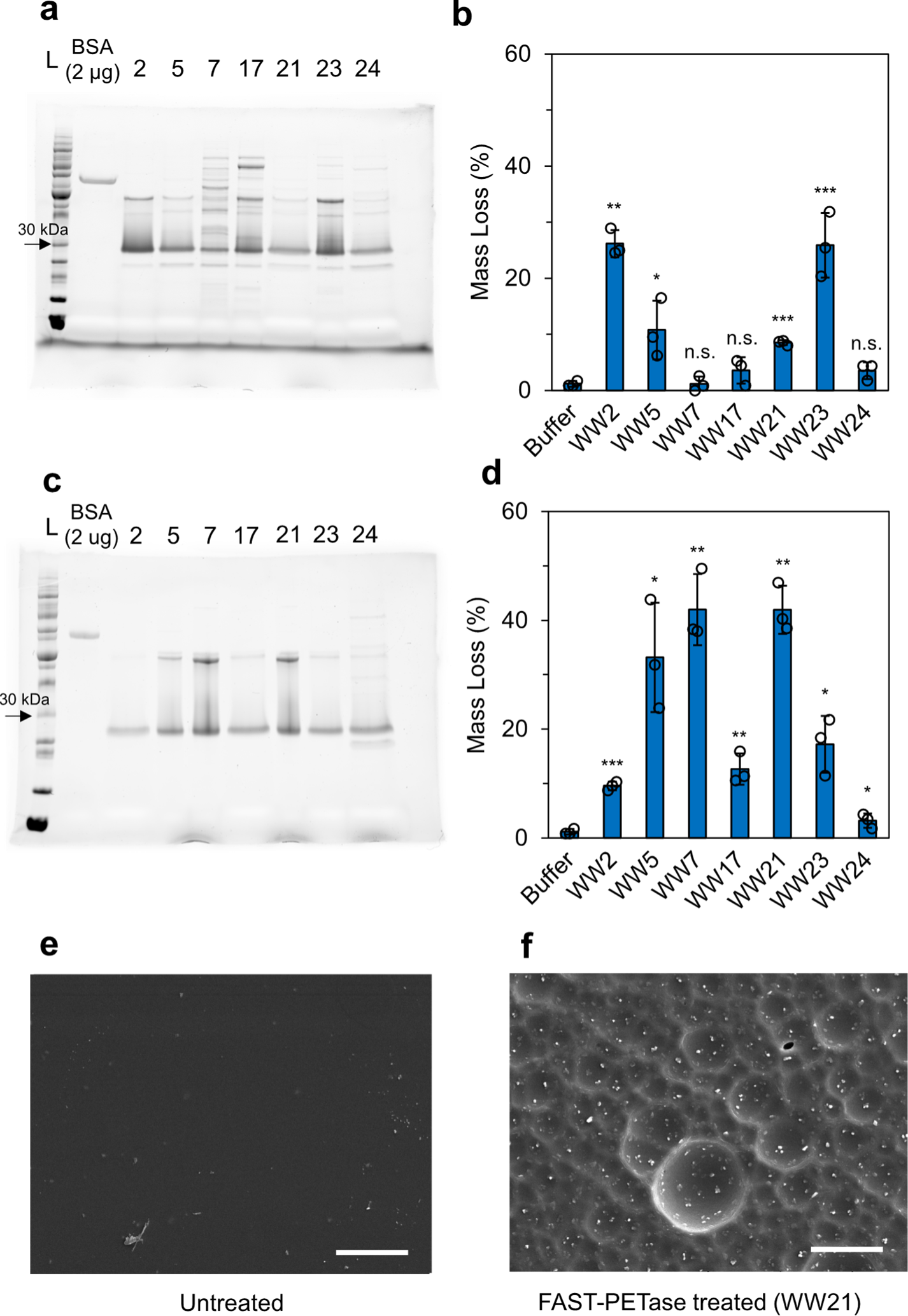
Enzymatic degradation of aPET films (7 mm diameter, 0.25 mm thick, ∼11 mg) by purified FAST-PETase from isolated transconjugants. SDS-PAGE gels of purified FAST-PETase are shown for **a**) cell lysates and **c**) culture supernatant. Mass loss from exposure to FAST-PETase purified from **b)** cell lysates and **d)** supernatant was measured after 96 h incubation at 50 °C. Bars show mean of three technical replicates for each isolate and control. Error bars: ± s.d. Statistical analyses were performed using Welch’s one-sided *t*-test to compare mass loss of each sample to the negative control (Buffer) (\**P* < 0.05, \*\**P* < 0.01, \*\*\**P* < 0.001). **e** SEM image of an untreated PET film. **f** SEM image of a PET film incubated at 50 °C for 96 h with FAST-PETase purified from WW21 culture supernatant. Scale bars: 50 μm.

### 2.4 Culture supernatant containing FAST-PETase degrades post-consumer PET plastic without enzymatic purification

To test if FAST-PETase produced from the isolates would degrade post-consumer PET without enzymatic purification, we incubated 6 mm hole punches of post-consumer PET (pcPET) from a coffee cup lid (crystallinity 11.4 ± 0.2% determined from differential scanning calorimetry, methods) with culture supernatant (M9 glycerol medium) diluted in 0.1 M KH_2_PO_4_ buffer. We made spectrophotometric measurements of degradation, which have been shown to be more sensitive than gravimetric measurements. The PET degradation byproducts, terephthalic acid (TPA) and mono-(2-hydroxyethyl)terephthalate (MHET), have both been shown to have an absorbance peak around 240 nm (A_240_) ^42^. In comparison to samples where FAST-PETase expression was either repressed or not induced, a significant increase in A_240_ occurred in samples exposed to supernatant from cultures where FAST-PETase expression was induced (Fig. 4a), indicating enzymatic degradation of the PET sample. Samples showing significant increase in A_240_ also showed signs of degradation observable with the naked eye (Fig. 4b). Further inspection by SEM showed roughening of the plastic surface of PET samples incubated with cultures induced to express FAST-PETase (Fig. 4b), which is characteristic of enzymatic degradation of PET ^13,21^.

**Figure 4.**
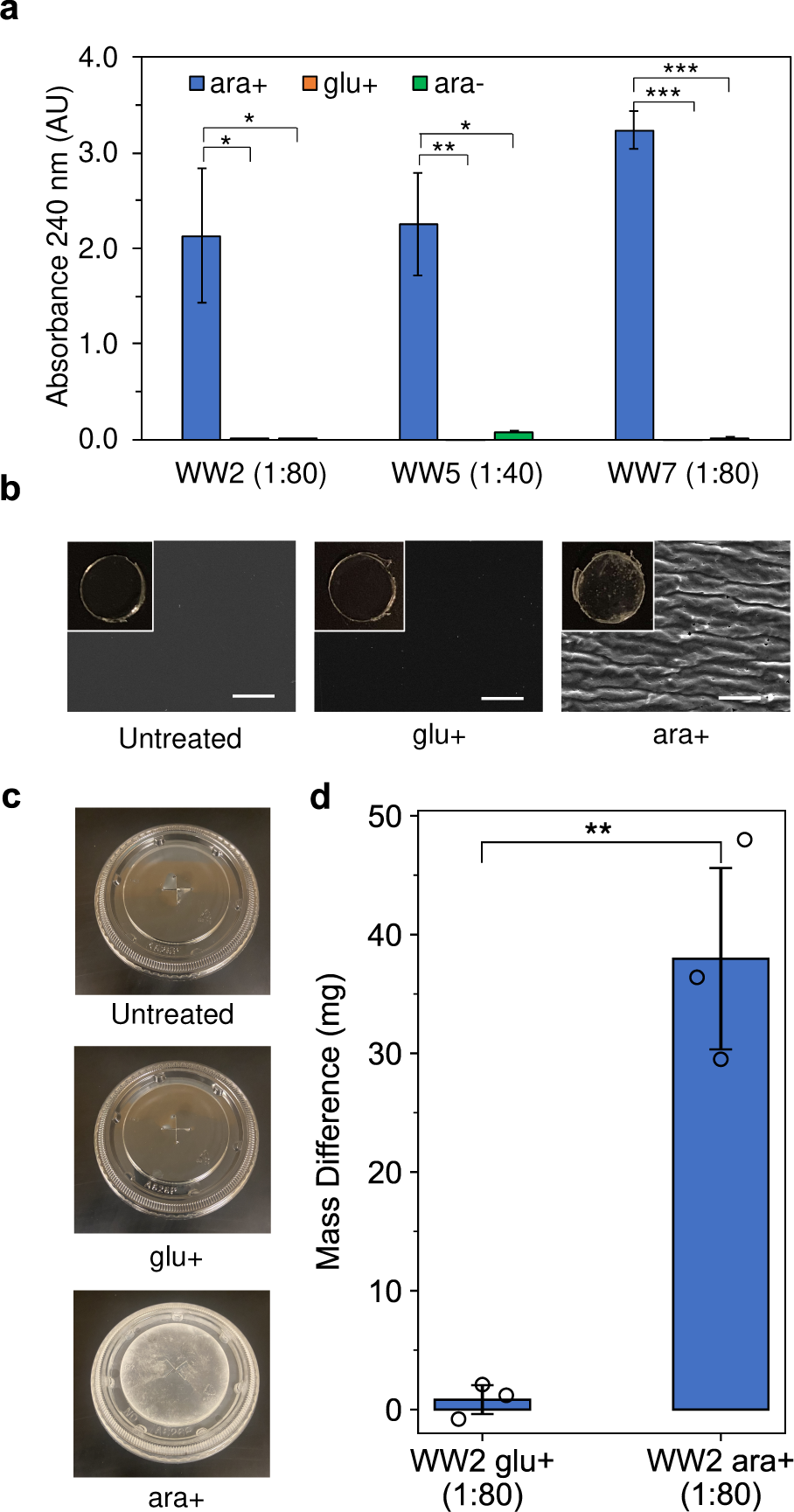
Degradation of a coffee cup lid using spent supernatant from engineered wastewater isolates diluted in 0.1 M KH_2_PO_4_-NaOH buffer (pH 8.0). **a** A_240_ measurements for diluted supernatant incubated with a 6 mm hole punch of pcPET for 120 h at 50 °C. ara+: protein expression induced with 0.2% arabinose; glu+: protein expression suppressed with 0.2% glucose; ara-: no arabinose added. Bars show mean of three technical replicates. Error bars: ± s.d. **b** Photos (inset) and SEM images of PET samples prior to treatment (left) or incubated in culture supernatant from WW5 (1:40) for 120 h. Scale bars: 50 μm. **c** Coffee cup lid (bottom) shows macroscopic signs of degradation after 168 h incubation at 50 °C with spent supernatant (ara+) from WW2 (1:80). An untreated lid (top) and lid incubated with spent supernatant (glu+) from WW2 (1:80) (middle) are shown for comparison. **d** Difference between initial and final mass of coffee cup lids incubated in supernatant from WW2 (1:80) after 168 h incubation at 50 °C. Bars show mean of three technical replicates. Error bars: ± s.d. All statistical analyses were performed using Welch’s one-sided *t*-test to compare ara+ samples to glu+ and ara-samples (\**P* < 0.05, \*\**P* < 0.01, \*\*\**P* < 0.001).

We hypothesized that higher enzyme concentration in undiluted culture supernatant would improve the PET degradation rate. We thus followed the same protocol with culture supernatant that had not been diluted in buffer. Contrary to our hypothesis, we observed minimal activity: there was little to no increase in A_240_ relative to media where FAST-PETase expression was suppressed (Supplementary Fig. 3). The same negative result was observed when we repeated this experiment with an additional pH adjustment step (addition of 1 M HCl) to modify the pH of the supernatant from 6.5 to 8.0, which is optimal for FAST-PETase activity ^21^ (Supplementary Fig. 3).

We tested scaling of pcPET degradation by increasing the total reaction volume, allowing an entire coffee cup lid (same pcPET product used in Fig. 4a) to incubate in diluted supernatant from WW2 (*Pusillimonas ssp.*) for 168 h at 50 °C. A lid that was incubated with supernatant where FAST-PETase expression had been induced showed signs of degradation visible to the naked eye (Fig. 4c) and significant mass loss (1.3 ± 0.3% of the initial mass of the lid; Fig. 4d). In comparison, a lid incubated with supernatant where FAST-PETase expression had been suppressed did not show any visible signs of degradation (Fig. 4c) nor appreciable mass loss (Fig. 4d).

## 3 Discussion

Engineering microbiomes to enable *in situ* degradation of plastics in diverse ecological niches remains a key milestone for deploying biological solutions for plastic waste at scale ^43^. Biotechnological plastic degradation and associated re-/upcycling can offer a means to achieve a circular life cycle for plastic waste in a sustainable and cost-effective manner, facilitating a broader move towards a circular economy. Developments in biotechnological approaches to plastic waste management will ideally complement related strategies for reduction of single-use plastics and innovations in biodegradable plastics.

In this work, we demonstrated a proof-of-concept for genetically engineering bacteria from municipal wastewater *in situ* to degrade PET plastics by using a broad-host-range, conjugating plasmid carrying the FAST-PETase gene. Previous studies have focused on optimizing expression and secretion of PET-degrading enzymes in single species ^44–46^, or employing naturally occurring and designer microbial communities to degrade PET plastics ^14,15,47,48^. In contrast, we genetically engineered bacterial isolates from an environmental sample to degrade PET via expression of an engineered PETase that exhibits high activity at moderate temperatures. Compared to cell augmentation efforts, *in situ* genetic bioaugmentation bypasses the adaptation barrier to a new environment, and thus could be used to achieve long-term bioremediation solutions. In particular, an approach focused on degradation of specific plastic compounds could be useful for removing microplastics from the waste streams of plastic manufacturing and recycling facilities. (A recent study reported that 6% of plastic processed at a recycling facility was found as microplastics in the facility’s wash water post-filtration ^49^.)

The approach demonstrated here may generalize to a variety of ecological niches: several studies have utilized the RK2/RP4 conjugation machinery coupled with a broad-host-range origin of replication (or a library of such origins) to engineer microbes from diverse environments: soil ^50^, mammalian guts ^51^, and marine ecosystems ^52^. A broad-host-range genetic engineering platform could facilitate construction of microbial consortia for degradation of individual types of plastic polymers that have known biodegradation pathways (e.g., PET). Such libraries of consortia could eventually be further expanded for degradation of other common plastic polymers, potentially resulting in modular tools for degradation of mixed-waste plastics.

Our results demonstrate that active FAST-PETase released by the engineered isolates can degrade PET without the need for enzymatic purification (Fig. 4). The assays using purified enzyme (Fig. 3) illustrate the plastic-degrading potential of the engineered isolates, which can guide future optimization efforts. In particular, our preliminary attempts to achieve degradation in LB medium showed no activity (isolation from WW5; data not shown). Moreover, we found that the solution conditions affect FAST-PETase activity: it was necessary to dilute the supernatant to achieve significant PET degradation, consistent with previous observations that FAST-PETase activity is inhibited at high enzyme concentrations ^53^ or by the presence of factors (e.g. other proteins and salts) in the supernatant. Further optimization of operating conditions could lead to significant improvements in enzyme activity in an environmental setting. Promoter choice for wide-spread or specific constitutive expression, optimization of ribosome-binding sites and transcriptional terminators, and screening of signal peptides in members of the microbiome of interest could likewise enhance the overall PET-degrading capacity ^54,55^. Furthermore, environmentally-triggered expression can minimize metabolic load and may be useful for avoiding unintended plastic degradation in e.g., product usage, storage, and infrastructure.

To achieve complete conversion of PET to its constituents TPA and ethylene glycol (EG), the PETase activity investigated here could be complemented by a MHETase enzyme, which would enable conversion of the depolymerization product, MHET, into TPA and EG ^13^. The turnover rate of wild-type MHETase from *Ideonella sakaiensis* (*k_cat_* ∼ 25 s^-1^ at 30 °C) ^38^ is much higher than that of FAST-PETase (*k_cat_* ∼ 0.016 s^-1^ at 50 °C) ^a^, so depolymerization by PETase will likely be the rate-limiting step for the conversion of PET into TPA and EG. Moreover, MHETase has been observed to improve PETase activity by relieving product inhibition caused by MHET monomers ^38,57^. Recently, a MHETase that is thermostable at the optimum operating temperature for FAST-PETase has been designed ^58^, offering an attractive dual-enzyme system for applications in PET biodegradation and upcycling. If production of large quantities of TPA and EG pose ecological or environmental concerns, bioaugmentation of additional consortia could be considered, using, for example, the consortium designed by Bao et al. ^59^ for efficient conversion of TPA and EG into biomass.

Using *E. coli* as a donor strain, we found that pFAST-PETase-cis could conjugate on a filter-mating setup and in native wastewater conditions at efficiencies high enough to allow isolation of transconjugants by plating. Improvements in conjugation efficiency could be achieved by exploring alternative donor strains, conjugation systems, and plasmid incompatibility classes. Moreover, the concentration of suspended solids (not measured) could affect conjugation efficiency by increasing the total surface area available for attachment, promoting spatial proximity between donor and recipient bacteria ^60^. We observed low taxonomic diversity among the isolates, likely in part due to our reliance on *mCherry* expression, driven by the P_bs_ promoter whose activity has only been previously reported in *Escherichia coli*, *Bacillus subtilis*, *and Saccharomyces cerevisiae* ^61^. Moreover, it is possible that conjugation efficiency from *E. coli* is higher towards closely-related species ^62^. Finally, the laboratory growth conditions likely had an impact on the results; reduced diversity of transconjugants has been reported in mating conditions that are less conducive to growth of diverse species ^51^. Additional improvements for selection of the plasmid could be made by using a promoter library for selection markers and origins of replication (e.g., ref 51).

We observed that pFAST-PETase-cis was lost in the absence of selection (Fig. 2), highlighting that long-term biodegradation applications will require efforts toward genetic stability. Improved maintenance of the function of interest could be achieved by incorporating plasmid maintenance systems such as toxin-antitoxin systems ^63,64^, gene entanglement ^65,66^, and plasmid partitioning systems ^67^ into the plasmid design, or by integrating the gene of interest into the chromosome of the host ^51^. Biocontainment systems to prevent spread of the plastic-degrading functionality to pathogenic bacteria should also be considered to avoid selection or amplification of pathogens.

Deployment of genetically modified microorganisms for degrading plastics will require careful consideration of environmental and public safety. To date, there have been few pilot-scale or field studies utilizing genetic bioaugmentation for bioremediation purposes ^34,68,69^. One notable case was a field release in Estonia in 1989 of *Pseudomonas putida* cultures carrying a gene for metabolizing phenol into a watershed contaminated from a phenol release caused by a subterranean oil shale mine fire ^70^. Six years after the release, the introduced operon was still found to be present in native microbiota ^70^. This study suggests that an introduced functionality could persist long-term in the environment if under selection for metabolism of an abundant pollutant. It will be prudent to test the use of engineered microbes for plastic degradation in contained systems while simultaneously assessing ecological risk, including: characterizing the modified the microorganisms and nature of the genetic modification, tracking the fate of these microorganisms and the introduced genes of interest, assessing the environmental impact of the release, and monitoring effects of the release on non-target microorganisms ^71^. The controlled environments in wastewater treatment units offer opportunities to assess ecological risks and explore performance of *in situ* microplastic degradation at scale.

## 4 Methods

### 4.1 Bacterial strains

*E. coli* DH5α (Matthew Scott Lab, University of Waterloo) and NEB10-β (New England Biolabs (NEB), Whitby, Ontario, Canada) were used for cloning steps. *E. coli* K-12 (Δ*ilvD*::*FRT*, Δ*galK*::*cfp-bla,* pSAS31, pTGD, pFAST-PETase-cis) was used as a conjugative donor strain for matings with wastewater samples. Engineered wastewater isolates (Table 1) were used for protein expression as indicated. Plasmid construction

All polymerase chain reaction (PCR) amplification steps were performed using Phusion DNA polymerase (NEB, Whitby, Ontario, Canada). Primers used for PCR amplification were ordered from Integrated DNA Technologies (IDT, Coralville, Iowa, United States) and are listed in Supplementary Data 3. PCR products were assembled using NEB Hifi DNA Assembly Master Mix (NEB) according to the manufacturer instructions. pNuc-trans-mCherry was constructed by replacing *mRFP* under the *P_bs_* promoter on pNuc-trans-mRFP ^60^ (gift from Thomas Hamilton and David Edgell, Western University) with *mCherry*. The backbone of pNuc-trans-mRFP was amplified using primers AY-9 and AY-10, and *mCherry* from pBT1-proD-mCherry (Addgene plasmid #65823, gift from Xiaoxia Lin, University of Michigan) using primers AY-7 and AY-8, followed by heat shock transformation into *E. coli* DH5α. pFAST-PETase-trans was assembled using PCR-amplified fragments from pNuc-trans-mCherry (primers OM-5 and OM-6), pFA6a-link-yoEGFP-SpHis5 (primers OM-8 and OM-9) (Addgene plasmid #44836), and pBTK522::FAST-PETase ^21^ (gift from Hal Alper, University of Texas at Austin) (primers OM-1 and OM-2), and was subsequently transformed into *E. coli* DH5α by heat shock. pFAST-PETase-cis was constructed by amplifying pFAST-PETase-trans (primers DE-3124 and DE-3125) ^60^ with 60-bp homologous overlaps to the AvrII cut site in pTA-Mob ^39^. The pTA-Mob plasmid (gift from Thomas Hamilton and David Edgell, Western University) was linearized using AvrII (NEB) and combined with the amplified fragment from pFAST-PETase-cis using a modified NEB Hifi Assembly protocol with a two-hour assembly step at 50 °C, followed by electroporation into electrocompetent *E. coli* NEB10-β.

### 4.2 Filter-mating conjugation and selection of engineered isolates

An auxotrophic donor strain was constructed by filter-mating *E. coli* NEB10-β (pFAST-PETase-cis) with *E. coli* K-12 (Δ*ilvD*::*FRT*, Δ*galK*::*cfp-bla,* pSAS31, pTGD) (gift from Xiaoxia Lin, University of Michigan) using a procedure similar to that described in ref. 72. Transconjugants were selected on Luria-Bertani (LB) plates (10 g/L tryptone, 10 g/L NaCl, 5 g/L yeast extract, 15 g/L agar) supplemented with ampicillin (100 μg/mL), gentamycin (50 μg/mL), and kanamycin (50 μg/mL), and were used as donors in filter-mating procedures with wastewater samples.

Untreated wastewater samples were collected from the City of Waterloo wastewater system access point (Wayne Parker lab, University of Waterloo, Ontario, Canada) and were processed the same day. 50 mL of wastewater was filtered through a sterile 0.40 μm-pore polycarbonate filter (Sigma-Aldrich, Oakville, Ontario, Canada), under vacuum. The filter and its contents were then vortexed in 1 mL of 1× phosphate-buffered saline (PBS; 2.56 g/L Na_2_HPO_4_·7H_2_O, 8 g/L NaCl, 0.2 g/L KCl, 0.2 g/L KH_2_PO_4_) for 1 min. Three washes, consisting of centrifuging for 5 min at 4000×*g* followed by resuspension in 1× PBS, were then completed. The solution was resuspended to an OD_600_ of 0.5. The donor strain, which was grown overnight from single colony in LB medium (10 g/L tryptone, 10 g/L NaCl, 5 g/L yeast extract) supplemented with appropriate antibiotics, was diluted 1:50 in LB with appropriate antibiotics, and was grown at 37 °C with shaking until exponential phase (OD_600_ of ∼0.5). The donor culture was then pelleted by centrifugation and resuspended to an OD of 0.5 in 1× PBS. Filter-mating of the donor culture and collected solids from the wastewater sample was completed by adding 100 μL of each onto a sterile 0.4 μm-pore polycarbonate filter placed on R2A agar (0.5 g/L yeast extract, 0.5 g/L proteose peptone no. 3, 0.5 g/L casamino acids, 0.5 g/L glucose, 0.5 g/L soluble starch, 0.3 g/L sodium pyruvate, 2.2 mM KH_2_PO_4_, 0.2 mM MgSO_4_, 15 g/L agar). Mating was allowed to occur for ∼24 hours before the filters were vortexed in 1 mL of 1× PBS for 1 min, and serial dilutions were plated on M9 glucose agar (2 g/L glucose, 2 mM MgSO_4_, 0.1 mM CaCl_2_, 64 g/L Na_2_HPO_4_·7H_2_O, 15 g/L KH_2_PO_4_, 2.5 g/L NaCl, 5.0 g/L NH_4_Cl, 15 g/L agar) supplemented with 50 µg/mL gentamycin and 100 µg/mL ampicillin. Plates were incubated at 30 °C until transconjugants appeared, which were observed after 1-2 days as red/pink colonies. Colonies that were identified as expressing *mCherry* based on fluorescence at 670 nm (Cy5 channel) using a Chemidoc MP Imaging System (Bio-Rad, Mississauga, Ontario, Canada) were streaked onto M9 glucose plates supplemented with 50 μg/mL gentamycin and 100 μg/mL ampicillin until pure cultures were obtained.

### 4.3 Conjugation efficiency in wastewater conditions

Untreated wastewater samples were collected from the City of Waterloo wastewater system access point (Wayne Parker lab, University of Waterloo) and were processed the same day. The donor strain *E. coli* K-12 (Δ*ilvD*::*FRT*, Δ*galK*::*cfp-bla,* pSAS31, pTGD, pFAST-PETase-cis) was grown from a single colony overnight in LB supplemented with 0.2% glucose and appropriate antibiotics, then diluted 1:50 into fresh LB with glucose and antibiotics. Once the donor strain reached an OD_600_ = 0.4-0.6, 50 mL of donor culture was washed twice in sterile 1× PBS and resuspended in 50 mL of sterile 1× PBS. Five mL of donor culture was added to either 20 mL of wastewater or 20 mL of wastewater with 10× concentrated solids (prepared by centrifuging 200 mL of wastewater at 2550×*g* for 15 min and decanting 180 mL of supernatant) in a 50 mL falcon tube, then incubated for 24 h at 30 °C with 60 rpm shaking. After 24 h, 1 mL of culture was washed twice in sterile 1× PBS. Serial dilutions of this mixture were then plated on selective M9 glucose plates containing gentamycin and ampicillin, and on non-selective M9 glucose plates. The plates were incubated at 30 °C for five days. Transconjugants were enumerated by counting colonies identified as expressing *mCherry* by visual inspection and by imaging under the Cy5 channel on a Chemidoc MP Imaging System. Total recipients were enumerated by counting colonies on non-selective plates. Conjugation efficiency was determined as the ratio of transconjugants to the total of recipients plus transconjugants.

### 4.4 Plasmid stability

A single colony was inoculated into 3 mL of LB media and grown at 30 °C with shaking. The culture was diluted 1:1000 into 3 mL of fresh growth medium once per day. At selected time points (Day 1, 3, 7, 10, 14), the culture was serially diluted in sterile 1× PBS and plated on selective (gentamycin at 50 μg/mL and ampicillin at 100 μg/mL) and non-selective LB plates. Plates were incubated at 30 °C for 16-24 h, and colony forming units were counted manually on each plate to quantify the cell density.

### 4.5 Protein expression and purification

A single colony was inoculated into LB supplemented with ampicillin (100 μg/mL) and gentamycin (50 μg/mL) and grown overnight at 30 °C with shaking. The overnight culture was then diluted 1:50 into 50 mL of LB supplemented with ampicillin and gentamycin. Protein expression was induced with 0.2% arabinose when the OD_600_ of the culture reached 0.6-0.8, after which the cells were cultured for ∼20 h at 30 °C with shaking. For purification of FAST-PETase from cell extracts, 50 mL of induced culture was centrifuged at 2550×*g* for 15 min at 4 °C and resuspended in 10 mL of cold lysis buffer (20 mM sodium phosphate, 300 mM NaCl, 10 mM imidazole, pH 7.4). The cells were incubated on ice with 0.1 mg/mL lysozyme for 30 min. The cells were further lysed by sonication in three 1 min intervals each followed by 1 min cool-down (Qsonica XL-2000, 3.2 mm, 8 W). Cell debris was pelleted at 2550×*g* for 30 min, and the 6×His-tagged FAST-PETase was purified from the supernatant by Ni-NTA affinity chromatography. 1 mL of 50% Ni-NTA agarose (Qiagen, Toronto, Ontario, Canada) was equilibrated using 10 bed volumes of wash buffer (same formulation as lysis buffer). The cell lysate was passed through the column, followed by washing twice with 10 bed volumes of wash buffer twice. The bound proteins were eluted using an imidazole step gradient of 50, 75, 100, and 250 mM in five elution steps using 2 bed volumes of elution buffer (20 mM sodium phosphate, 300 mM NaCl, pH 7.4. Imidazole concentrations in each elution step were 50, 75, 100, 250, 250 mM, respectively). For purification of FAST-PETase from culture supernatant, 50 mL of induced culture was centrifuged at 2550×*g* and 4 °C for 30 min, and FAST-PETase was purified from the supernatant using the same chromatography procedure described above.

Protein purity for each elution was checked by SDS-PAGE using stain-free imaging on a Chemidoc MP Imaging System. The purified protein from elutions containing the fewest contaminating proteins (typically elution steps 3-5) were concentrated in 1× PBS using a 5 kDa cut-off Vivaspin Centrifugal Concentrator (Sartorius, Oakville, Ontario, Canada). Protein concentrations were determined using the micro-BCA assay (Thermo Fisher Scientific, Mississauga, Ontario, Canada), where protein concentrations were correlated to bovine serum albumin standards at an absorbance of 562 nm in a plate reader.

### 4.6 PET sample preparation

Amorphous PET (aPET) film from GoodFellow (Pittsburgh, Pennsylvania, United States; product ES301445; specification: 0.25 mm thick, 1.300-1.400 g cm^-3^ density, 1.580-1.640 refractive index, 100×10^-13^ cm^3^ *⋅* cm cm^-2^ s^-1^ Pa^-1^ permeability to water at 25 °C) and post-consumer PET (pcPET) (coffee cup lid No. A626P, Amhil North America, Mississauga, Ontario, Canada) were prepared in circular form (aPET: 7 mm diameter, pcPET: 6 mm diameter) using office hole punchers. The PET discs were washed for 30 min with 1% SDS, 20% ethanol, and distilled water, and dried overnight at 50 °C. Prior to degradation experiments, individual PET discs were weighed on an analytical balance with accuracy of ± 0.1 mg.

### 4.7 PET degradation assay with purified FAST-PETase

A single, pre-weighed aPET disc was placed in a 2 mL polypropylene microcentrifuge tube with 600 μL of 0.1 M KH_2_PO_4_-NaOH buffer (pH 8.0) with 3.6 μg of purified protein concentrate (corresponding to 200 nM FAST-PETase). The tube was capped to prevent volatilization. The sample was incubated at 50 °C for 96 h, replenishing with fresh buffer and enzyme solution every 24 h to maximize the degradation rate. The PET samples were subsequently washed with 1% SDS, 20% ethanol, and distilled water, dried for 24 h at 50 °C, then weighed on an analytical balance to determine weight loss.

### 4.8 PET degradation with culture supernatant

A single colony was inoculated into 10 mL of M9 glycerol (4 g/L glycerol, 2 mM MgSO_4_, 0.1 mM CaCl_2_, 64 g/L Na_2_HPO_4_·7H_2_O, 15 g/L KH_2_PO_4_, 2.5 g/L NaCl, 5.0 g/L NH_4_Cl) supplemented with ampicillin (100 μg/mL) and gentamycin (100 μg/mL) and grown until saturation at 30 °C with shaking. The saturated culture was then diluted 1:50 into 50 mL of M9 glycerol supplemented with ampicillin and gentamycin. Protein expression was induced with 0.2% arabinose, not induced, or repressed with 0.2% glucose when the OD_600_ of the culture reached 0.3-0.5, after which the cells were cultured for 12-14 h at 30 °C with shaking (220 rpm). The supernatant for each culture was collected by centrifuging 50 mL of culture at 2550×*g* for 60 min at 4 °C and stored for the duration of the experiment at 4 °C. A single pcPET disc was placed in a 2 mL polypropylene microcentrifuge tube with 2 mL of supernatant. A replicate tube was prepared without PET for comparison to account for any change in absorbance from other compounds. All samples were incubated at 50 °C for 120 h. After incubation, samples were centrifuged at 14000×*g* for 3 min. The supernatant was transferred to a quartz cuvette, and absorbance at 240 nm was measured using UV-spectrophotometry ^42,73^, where the spectrophotometer was blanked to 0.1 M KH_2_PO_4_-NaOH buffer (pH 8). The mean absorbance of samples without PET was subtracted from the absorbance reading of samples that contained PET.

For degradation of whole coffee cup lids, the lids were washed, dried, and pre-weighed using the same procedure described in Section 4.6. The lids were submerged in individual sealed containers with 150 mL of 0.1 M KH_2_PO_4_-NaOH buffer (pH 8) containing 1:80 culture supernatant from WW2. The reaction was carried out for 168 h at 50 °C. The final mass of the lids were measured after washing and drying using the same procedure described in Section 4.6.

### 4.9 Scanning electron microscopy

PET samples were washed with 1% SDS, 20% ethanol, and distilled water, then dried overnight at 50 °C. The samples were attached to an aluminum stub using double-sided carbon tape, followed by sputter-coating (Polaron Instruments SEM Coating Unit E5100) with gold particles for 2 min at 20 mA in an argon atmosphere (layer thickness ∼30 nm). The samples were then imaged by scanning electron microscopy (Tescan VEGA TS-5130) at 20 kV and 1000× magnification.

### 4.10 Differential scanning calorimetry

PET film crystallinity was determined using differential scanning calorimetry. PET film samples (9-11 mg) were loaded into aluminum Tzero pans (TA Instruments, Grimsby, Ontario, Canada; product 901683.901) with a Tzero lid (TA instruments; product 901671.901) for solid samples. Samples were heated from 40 to 300 °C at 10 °C min^-1^, held at 300 °C for 1 min, cooled from 300 to 30 °C at 10 °C min^−1^, and held at 30 °C for 1 min in a Q2000 calorimeter (TA Instruments) with an RCS90 refrigerated cooling system. Percentage crystallinity was calculated as:

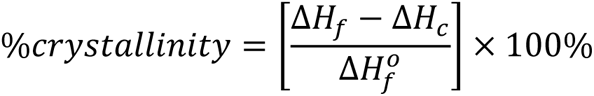

where *ΔH_f_* is the enthalpy of fusion at the melting point, *ΔH_c_* is the enthalpy of crystallization, and 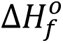 is the enthalpy of fusion for a completely crystalline sample at its equilibrium melting temperature, which is 140 J g^-1^ (ref. 74). TA Universal Analysis software was used to calculate *ΔH_f_* and *ΔH_c_*. *ΔH_f_* and *ΔH_c_* were measured by integrating from 110-120 °C to 150 °C and 210 °C to 260 °C, respectively, with a linear baseline. Crystallinity was measured in triplicate for each source of PET.

### 4.11 Statistical analysis

Experiments were performed in biological and technical replicates where mentioned in figure captions. All statistical analyses were performed in Python 3.9 using the SciPy ^75^ library (v1.7.3). Welch’s one-sided *t*-test was used to compare the means of two groups. *P* < 0.05 was considered as statistically significant.

## Acknowledgments

This work was supported by funding from the Natural Sciences and Engineering Research Council of Canada (NSERC): RGPIN-03826-2018 (B.P.I.); 355513-2017 (M.G.A.); CGS-D (A.Y.). A.Y. was also supported by funding from the Ken O’Driscoll Graduate Scholarship in Polymer Engineering/Science, and Murray Moo Young Biochemical Engineering Scholarship. We are thankful to our lab assistants for their support in preparing materials and reagents (Melissa Meighoo, Julie Kim, Joyce Yang), cloning procedures (Melissa Meighoo), and CFU counts (Nitya Yadav). We thank Narasimman Lakshminarasimman and Wayne Parker (University of Waterloo) for providing wastewater samples. We thank Michael Lynch (Metagenom Bio Life Science) for providing information about the bITS sequencing protocol. This work was carried out on the Haldimand Tract, the land granted to the Six Nations that includes six miles on each side of the Grand River.

## Contributions

A.Y., M.G.A., and B.P.I conceptualized the research. A.Y., O.D.M., M.G.A., and B.P.I. conceived the experiments. A.Y. and O.D.M. designed and constructed plasmids and performed filter-matings. A.Y., O.D.M. and K.C.H. performed protein purifications, prepared PET samples, and performed PET degradation assays. A.Y. performed conjugation experiments in native wastewater conditions and plasmid stability experiments. A.Y., M.G.A., and B.P.I. analyzed the results. M.G.A. and B.P.I. provided supervision and resources for this study. A.Y., O.D.M., M.G.A., and B.P.I. wrote the original draft of the manuscript. A.Y., M.G.A., and B.P.I. revised the manuscript.

## Footnotes

a Converted from the specific activity of 348 μmol_TPAeq_ h^-1^ mg_enzyme_^-1 56^.

