## Supplementary Information for "Degradation of PET Plastics by Wastewater Bacteria Engineered via Conjugation"

### Supplementary Methods

#### 1.1 Identification of Isolates

Pure cultures were submitted to Metagenom Bio Life Science (Waterloo, Ontario, Canada) for identification by sequencing the bacterial internal transcribed spacer (ITS) region [1]. The company provided the following details about their protocol: "Genomic DNA was extracted using the Metagenom Bio Sox DNA Isolation Kit (No. 18011-50). Full length PCR primers as described below were designed and synthesized by IDT. Illumina adapter and 8-nt index sequences were added at the 5' ends of bacterial ITS forward [UNI_ITS_fw: 5' KRGGRYKAAGTCGTAACAAG] and reverse primers [bacITS189r: 5' TACTDAGATGTTTCASTTC] respectively [1,2]. In order to increase $T_m$, CGC and CGCG were added to the 5' ends of forward and reverse primers respectively. PCR was set up in duplicate for each DNA sample (25 µl each). Reaction mixture consisted of 12.5 µl of Q5 High-Fidelity 2x Master Mix (NEB), 1.25 µl of 10 µM PCR forward primer, 1.25 µl of 10 µM PCR reverse primer, 5.0 µl eDNA, and 5.0 µl of PCR water. DNA was initially denatured at 98 °C for 3 min, followed by 35 cycles of 98 °C for 10 sec, 53 °C for 30 sec and 72 °C for 1 min, and then extended at 72 °C for 5 min. Duplicate PCR products were pooled and confirmed by 1.5% TAE agarose gel.

Remaining product was purified byAMPure XP beads and quantified using the Qubit dsDNA assay kit. Average sizes of bITS amplicons were estimated based on the major band on the gel. Amplicons were sequenced with MiSeq Reagent v2 kit (2x250 cycles), and FASTQ data were generated. The 3'ends of 16S rRNA are in MiSeq read 1, and the 5' ends of 23S sequences are in Read 2 respectively. Primers were removed from raw reads using cutadapt v.3.5 [3]. Reads were then quality filtered using DADA2 v.1.22 [4] with default parameters and removing all sequences with ambiguous nucleotides (N). Chimeric sequences were removed using the *dada2::removeBimeraDenovo()* method and due to the amplicon length the forward and reverse reads were processed separately and stored in separate amplified sequence variant (ASV) tables. ASVs constructed from forward reads were used in taxonomic assignment. Classification was performed using a naive Bayesian classifier implemented in DADA2, *dada2::assignTaxonomy()*, trained against the bacterial ITS region extracted from genomes in the Genome Taxonomy Database (GTDB) release 202 [5]. Species-level taxonomy was corrected by exact sequence classification using *dada2::assignSpecies()*. All analysis was performed using R 4.1.2."

#### 1.2 Estimation of FAST-PETase Concentrations from SDS-PAGE

Samples were mixed 1:1 with 2× sample loading dye (final concentration of 62.5 mM Tris-HCl pH 6.8, 2% SDS, 0.002% bromophenol blue, 5% β-mercaptoethanol, 10% glycerol) and boiled for 10 min at 95 °C. A total mass of 3.6 μg of purified protein for each sample (based on concentration measurements from the micro-BCA assay) was visualized using stain-free imaging of a stain-free acrylamide gel. The averaged pixel intensity of a 2 μg sample of bovine serum albumin (BSA) standard was used to standardize pixel intensity versus protein mass (9431 AU/μg for cell extracts, 6243 AU/μg for supernatant). The average intensity of each sample was then converted to mass (Supplementary Table 1 and Supplementary Table 2).

#### 1.3 Imaging Plates

Plates were imaged on a Chemidoc MP Imaging System (Bio-Rad) on the Cy5 and Colorimetric channels with an exposure time of 5 ms and 0.035 ms, respectively. 16-bit images for each channel were overlayed using ImageJ and the minimum displayed pixel intensity for each channel was adjusted to enhance viewing (Cy5: 9994; Colorimetric: 60026).

#### 1.4 Confirming auxotrophy of *E. coli* donor strain

An overnight culture grown in LB supplemented with ampicillin (100 μg/mL), gentamycin (50 ug/mL) and kanamycin (50 μg/mL) from a single colony and cultured at 37 °C with shaking. The overnight culture was diluted 1:50 into LB supplemented with the same antibiotics and was grown to early exponential phase ($OD_{600}$ ~ 0.5). The culture was washed twice in sterile 1× PBS and was resuspended in 1 mL of 1× PBS. 100 μL of culture was plated on M9 glucose agar supplemented with ampicillin (100 μg/mL), gentamycin (50 ug/mL) and kanamycin (50 μg/mL) and incubated at 30 °C for up to six days.

### Supplementary Tables

Supplementary Table 1. Estimated mass of FAST-PETase utilized in PET degradation assays from cell extracts (Fig. 3a).

| Isolate | Average Band Intensity (AU) | Mass of FAST-PETase (μg) |
|---|---|---|
| WW2 | 18862 | 4.6 |
| WW5 | 43150 | 3.5 |
| WW7 | 32829 | 3.2 |
| WW17 | 30341 | 4.1 |
| WW21 | 38962 | 2.5 |
| WW23 | 23712 | 4.3 |
| WW24 | 40447 | 2.6 |

Supplementary Table 2. Estimated mass of FAST-PETase utilized in PET depolymerization assays from supernatant (Fig. 3b).

| Isolate | Average Band Intensity (AU) | Mass of FAST-PETase (μg) |
|---|---|---|
| WW2 | 12486 | 2.0 |
| WW5 | 12205 | 3.9 |
| WW7 | 24403 | 5.1 |
| WW17 | 32010 | 3.1 |
| WW21 | 19432 | 4.9 |
| WW23 | 30765 | 3.3 |
| WW24 | 20344 | 3.3 |

### Supplementary Figures

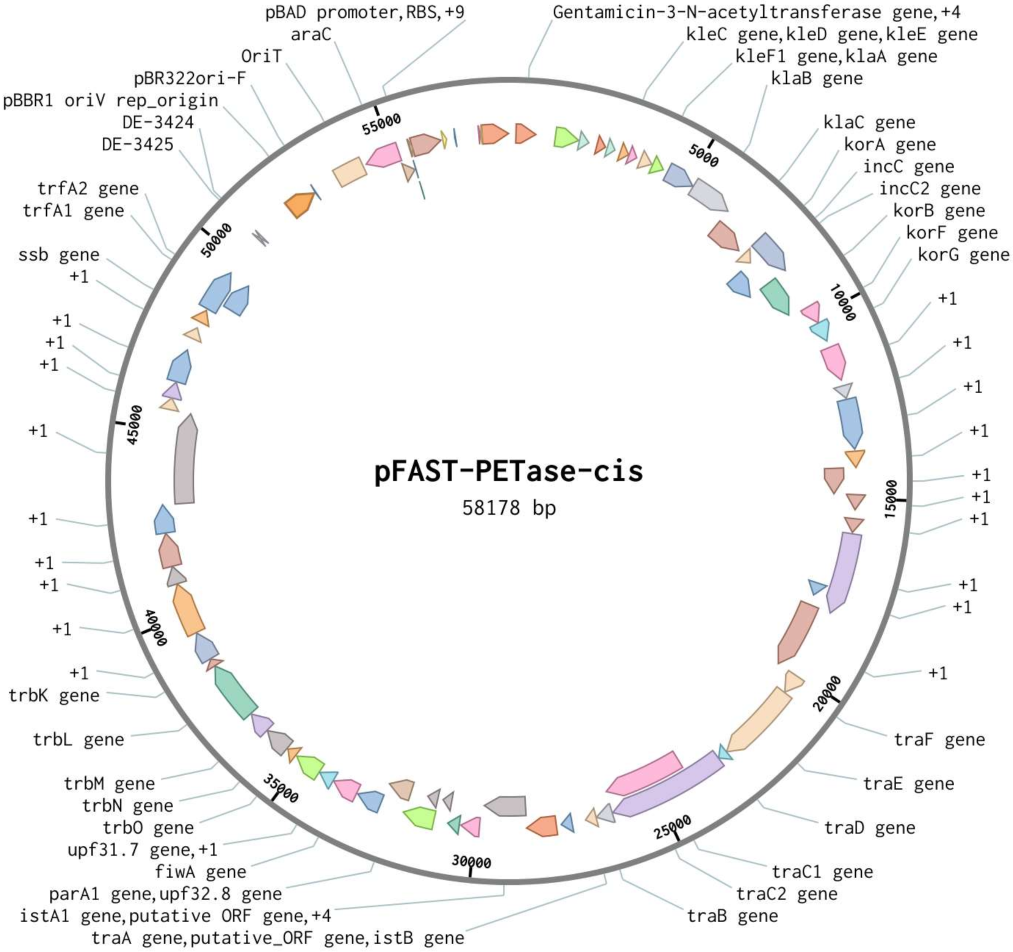


Supplementary Figure 1. Plasmid map of pFAST-PETase-cis. The corresponding GenBank formatted file of the nucleotide sequence is provided in Supplementary Data 4.

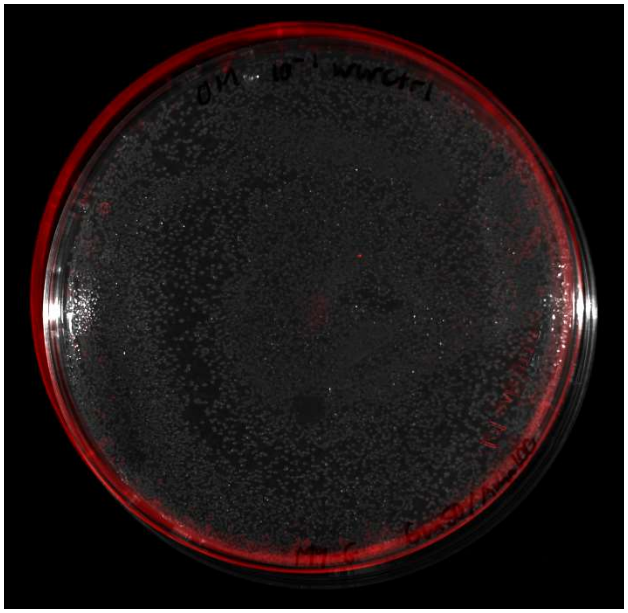

Supplementary Figure 2. Selection plate for wastewater sample without donor strain added (diluted 10-fold).

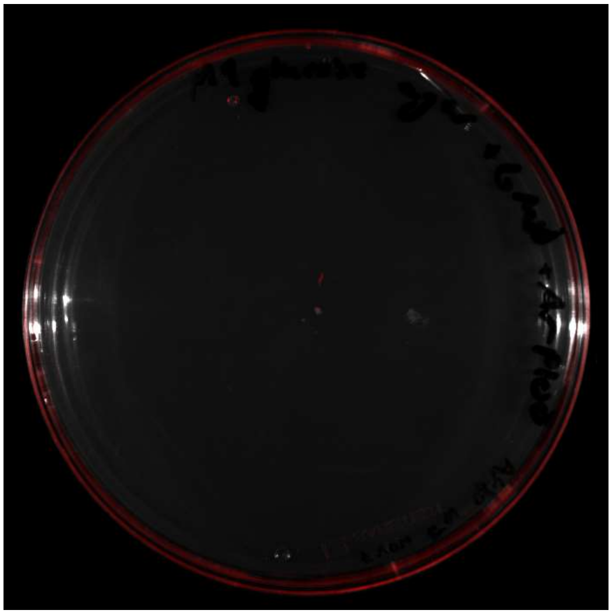

Supplementary Figure 3. Donor strain incubated at 30 °C on M9 glucose plate with antibiotics for 6 days.

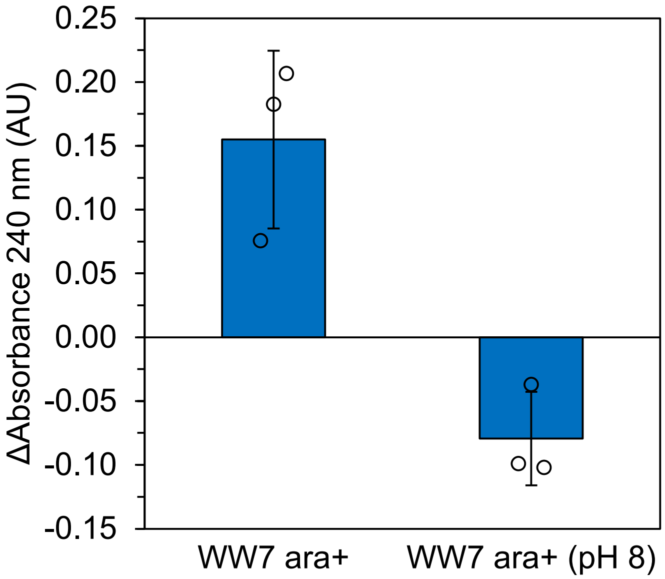


Supplementary Figure 4. $A_{240}$ measurements for M9 glycerol supernatant incubated with pcPET for 120 h at 50 °C. ara+: protein expression induced with 0.2% arabinose. Bars show mean of three technical replicates. Error bars: ± s.d.
